# Addressing differentiation in live human keratinocytes by assessment of membrane packing order

**DOI:** 10.1101/2020.06.14.151233

**Authors:** Danuta Gutowska-Owsiak, Christian Eggeling, Graham S Ogg, Jorge Bernardino de la Serna

## Abstract

Differentiation of keratinocytes is critical for epidermal stratification and formation of a protective stratum corneum. It involves a series of complex processes leading through gradual changes in characteristics and functions of keratinocytes up to their programmed cell death via cornification. The stratum corneum is an impermeable barrier, comprised of dead cell remnants (corneocytes) embedded within lipid matrix. Corneocyte membranes are comprised of specialized lipids linked to late differentiation proteins, contributing to the formation of a highly stiff and mechanically strengthen layer. To date, the assessment of the progression of keratinocyte differentiation is only possible by determination of specific differentiation markers, e.g. by using proteomics-based approaches. Unfortunately, this requires fixation or cell lysis, and currently there is no robust methodology available to study differentiation in living cells, neither at a single cell, nor in high throughput. Here, we explore a new live-cell based approaches for screening differentiation advancement in keratinocytes, in a “calcium switch” model. We employ a polarity-sensitive dye, Laurdan, and Laurdan general polarization function (GP) as a reporter of the degree of membrane lateral packing order or condensation, as an adequate marker of differentiation. We show that the assay is straightforward and can be conducted either on a single cell level using confocal spectral imaging or on the ensemble level using a fluorescence plate reader. Such systematic quantification may become useful for understanding mechanisms of keratinocyte differentiation, such as the role of membrane inhomogeneities in stiffness, and for future therapeutic development.

## Introduction

The epidermis, in the form of a stratified and desquamating tissue is a fascinating feature of many organisms; it retains very distinct characteristics which enable it to exert function, shielding us from external threats and preventing fluid evaporation. More and more is known on how these unique features contribute to the important protective role of the epidermis; these lessons are learned both from a physiological setting and through the perspective of skin diseases. For example, keratinocyte differentiation and cornification are essential for prevention of allergic diathesis by forming a functional skin barrier (1–4). Cornification encompasses a series of processes leading to programmed keratinocyte death, which results in formation of cellular remnant known as “corneocyte”. Corneocytes, embedded in a lipid-enriched matrix (5) provide mechanical hardness, while the lipids support elastic properties of the stratum corneum. In addition, at the cellular level, the lamellar organization of lipids and their lateral packing properties prevent evaporation (6–8), realizing a watertight seal over the moisture trapped in the stratum corneum. This phenomenon is achieved by high abundance of hydrophilic compounds, known as Natural Moisturising Factor (NMF).

Multiple processes take place during the cornification advancement in the epidermis; these occur simultaneously both at the organellar and molecular levels (Figure 1). The characteristics of progression in keratinocyte differentiation in vitro are reflected by multiple morphological changes (i.e. related to the cell shape and adherence) and distinct organellar transformations. For instance, the appearance and maturation of keratohyalin granules (KHGs) (9), nuclear condensation and extrusion (10, 11), cytoskeleton collapse (9), and alterations in mitochondria (12); in some of those events, autophagy has been shown to be involved (13). At the molecular level, shift in cytokeratin expression has been documented (keratin 5, keratin 14 are progressively replaced with keratin 1 and keratin 10) while additional proteins are upregulated, e.g. involucrin and transglutaminase as well as the late differentiation markers (filaggrin, loricrin) (5). These differentiation-dependent proteins contribute to barrier strengthening by incorporation into insoluble cornified envelope (CE; also known as cornified cell envelope, CCE) or by supporting the hydration of the stratum corneum (14, 15).

**Fig. 1.**
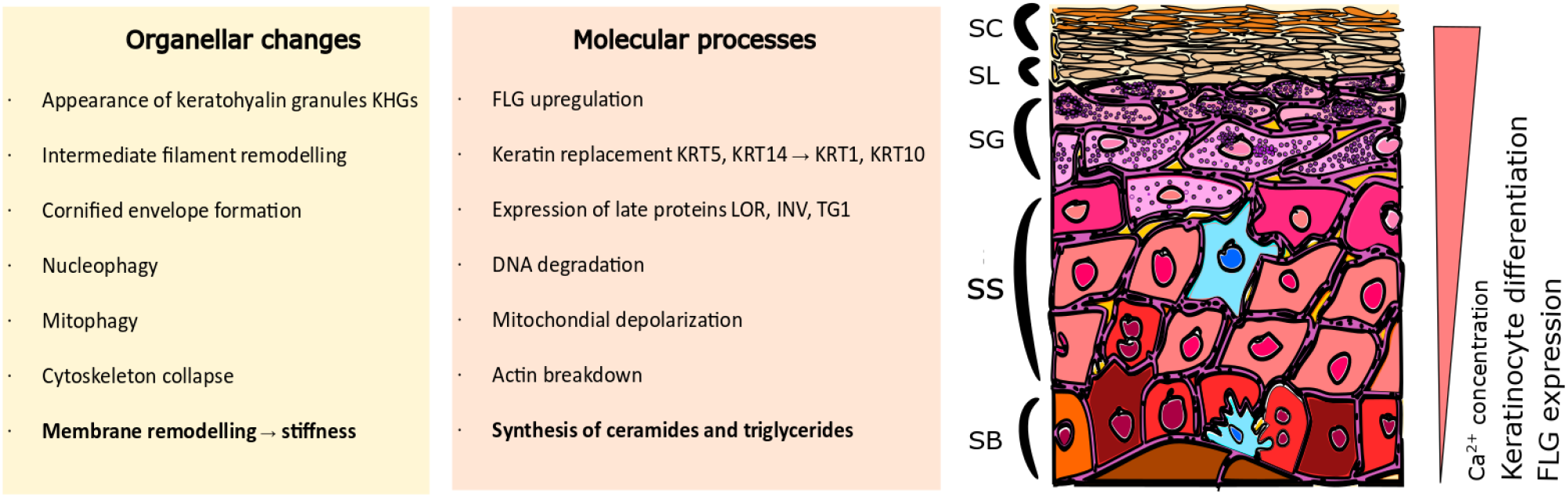
Scheme and Cartoon describing the organellar and molecular changes in keratinocytes during differentiation process. Stratified epidermis consists of multiple layers of epidermal keratinocytes characterised by increasing advancement of cell differentiation. Cells undergo extensive organellar and molecular alterations as they gradually move to the upper layers. SC: Stratum Corneum; SL: Stratum Lucidum; SG: Stratum Granulosum; SS: Stratum Spinosum; SB: Stratum Basale.

Along with the expression of protein markers and organellar changes, overall increase in cell stiffness can be observed (8, 16, 17). This stiffness is a consequence of two main processes; the formation of a rigid, protein-based CE and extensive lipid remodeling leading to specialized lipid profile adaptation (18–20). Specifically, the high phospholipid and cholesterol content is gradually replaced by a predominance of ceramides and saturated triglycerides (21), espe cially in conditions where stratification is supported. These lipid composition variations lead to more ordered membrane packing (8), higher stiffness and consequently, changes in the relative lateral heterogeneity properties of the plasma membrane (22).This selection has specific functional properties; the ceramides and free fatty acids provide attachment points for the CE proteins (18, 23, 24), and a scaffold to replace the plasma membrane by a corneocyte lipid envelope (CLE), which links CE with the lipids of the intercellular matrix (25). It is believed that ceramide enrichment displaces the balance between ordered and disordered regions to support membrane heterogeneity, where higher order regions work as docking molecular anchors supporting the CLE function and ultimately, stabilizing the CE structure. Regulation of ceramide synthesis is mediated via peroxisome proliferator-activated receptor (PPAR) pathways (26) associated with keratinocyte differentiation (27); disturbances in these pathways result in clinical manifestations (24).

To date, there is not available a robust experimental readout for the assessment of differentiation advancement in live primary keratinocytes. Protein-targeted detection methods, such as expression quantification by proteomics-based approaches, as well as other most commonly used, such as immunostaining or western blot, require cell permeabilization or lysis and are only suitable for end-point measurements.

Here, we built upon the known lipid profile variation and ceramide enrichment during keratinocyte differentiation; we hypothesised that lipid lateral packing and order increase can be used as a hallmark to quantitatively asses live-cell cornification advancement. For this purpose, we took advantage of the lipophilic polarity-sensitive membrane dye Laurdan (figure 2A, B), purposely designed by Weber and Farris (28) to have an electron-donor and electron-acceptor, which displays large solvent-dependent fluorescence shifts. Laurdan ubiquitously distributes at the plane of the plasma membrane, regardless of its lipid lateral packing properties, and is oriented parallel with the phospholipid’s hydrophobic tails in membranes (Figure 2B). Moreover, its emission spectra and location are independent of phospholipid head groups and notably, indirectly report the packing order by sensing the degree of water penetration it its vicinity (29–33). Most of these environmentally-sensitive probes display an increase in charge separation when excited in polar solvents resulting in a larger dipolar moment (34); the penetration of water molecules into a bilayer formed of loosely packed lipids allow more rotational and translational freedom to the probe, yielding an emission shift towards longer wavelengths. On the contrary, the probe emission shifts towards the blue when its motion is more restricted due to sensing a milieu with lesser water content (Figure 2C-D). Ratiometrically measuring the emission intensities at 440 and 490nm renders the Laurdan “general polarization index” (GP), which allows relative quantification of the membrane order or degree of lipid lateral packing (30, 31, 35). This method has been used in visualizing native membrane microdomains in planar supported bilayers, giant unilamellar vesicles (36, 37) and in living cells (9, 38–41).

**Fig. 2.**
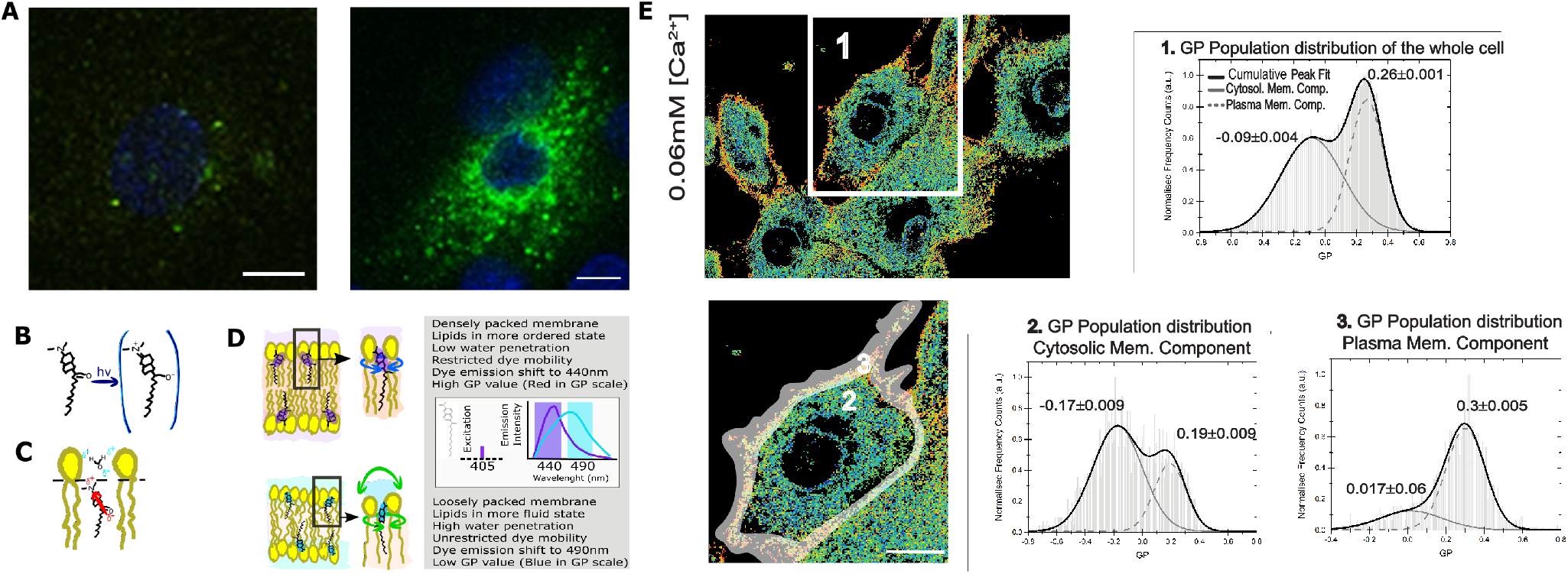
Membrane order assessment in live normal human epithelial keratinocytes (NHEKs) stiffness assessment in live cells by LAURDAN polarisation sensing dye. Membrane order assessment in live normal human epithelial keratinocytes (NHEKs) stiffness assessment in live cells by LAURDAN polarisation sensing dye. A) Detection of filaggrin-containing keratohyalin granules (KHGs) induced in primary normal human epidermal keratinocytes upon calcium switch by immunofluorescence (left panel represents [Ca^2 +^]=0.06 mM; right panel represents [Ca^2 +^]=1.5 mM)); scale bar 10µm; B) C) D) Cartoon representing Laurdan chemical structure, its positioning within membranes and a description of its biophysical and fluorescence spectral properties. Loosely packed lipids and high water content promote unrestricted Laurdan mobility, resulting in dye emission shift representing fluid membrane state while dense lipid packing and lower water penetration restrict free movement of the dye, resulting in reflecting of shorter wavelengths; E) The differences in LAURDAN emission reflects increased ordering of the lipids and is detected by a shift in emission wavelength. This is reflected in an increase in the value of General Polarization (GP). Pseudo-colour scale represents the increase as a gradual shift from fluid (blue) to stiff (red) membrane. Representative GP images as determined from 2D scanning confocal spectral imaging on live undifferentiated NHEKS [Ca^2 +^]=0.06mM): overview (upper panel, scale bar 5µm) and zoom-in (lower panel, grey-shaded area: selection for plasma membrane part) of area marked by a white box. Higher GP values (colour-coded from blue to red as labelled) indicate stiffer membranes. Frequency histograms of GP pixel values from areas of images marked by the respective number 1-3: 1) averaging over a whole cell; 2) cytosolic membranes fraction; and 3) plasma membrane fraction. The histograms were fit by a double-Gaussian (black line, with peak values and their accuracy from the fit stated) to outline the plasma membrane (right peaks) and cytosolic (left peaks) fraction. Analysis from 3 different cell batches, and a minimum of 5 cells per batch.

Even though the membrane order can only be measured indirectly, and being aware that the stiffness of a membrane influences can be in principle better characterised by fluorescence polarisation or fluorescence anisotropy, in this paper we introduced membrane lateral packing or order as a readout for keratinocyte differentiation advancement. Using single-cell based confocal spectral imaging or a fluorescence plate reader for high-throughput screening of cellular ensembles we utilize the Laurdan general polarisation index assessment of membrane order as an efficient readout for determination of the progression of keratinocyte differentiation in living cells.

## Material and Methods

### A. Human keratinocyte culture and calcium switch

Normal human epidermal keratinocytes (NHEKs) were purchased from Lonza (Lonza, Basel, Switzerland, neonatal, pooled) and cultured in keratinocyte KBM-2 media (Lonza, Basel, Switzerland) at conditions supporting proliferation ([Ca^2+^]=0.06mM) and subcultured by accutase treatment (Sigma Aldrich, Dorset, UK). To promote terminal differentiation, calcium switch was performed by adjusting calcium level do the final concentration of [Ca^2+^]=1.5mM for a pe riod of 24h before immunofluorescent labelling. To adjust to the desired calcium concentration a calcium switch was conducted over a period of 24h by replacing the culture media with media adjusted to the desired calcium concentration by adding CaCl_2_ (Sigma Aldrich, Dorset, UK sigma.)

### B. Fluorescent antibody staining and confocal microscopy imaging

NHEKs were grown and calcium-switched in eight-well cell culture slides (Beckton Dickinson), fixed and permeabilized by neat acetone. Blocking was carried out in freshly made blocking buffer (5% FCS, Sigma-Aldrich, Gillingham, Dorset, UK; 2% BSA, Sigma-Aldrich, Gillingham, Dorset, UK in PBS). Anti-filaggrin primary antibody staining (mouse monoclonal 15C10 from Leica Biosystems, Milton Keynes, UK) was followed by secondary antibody labelling (anti-mouse Alexa 488, Life Technologies/ThermoFisher Scientific, Waltham, MA, USA), all carried out in PBS. To visualize nuclei NucBlue reagent (Hoechst, Life Technologies/ThermoFisher Scientific, Waltham, MA, USA) was used. The cover-slides were mounted with Mowiol-488 (Sigma-Aldrich, Gillingham, Dorset, UK). Imaging acquisition was carried out on a Zeiss 780 inverted confocal microscope (Zeiss, Jena, Germany), by recording 2D images or 3D z-stacks. We used 488nm excitation line and detected from 500-550nm. Images were postprocessed using Zen Software (Zeiss, Jena, Germany) and ImageJ National Institutes of Health, Bethesda, Maryland, USA).

### C. Quantification of membrane order by spectral imaging

Spectral imaging of the different membrane samples was performed on a Zeiss LSM 780 confocal microscope equipped with a 32-channel GaAsP detector array. Excitation laser at 405nm was used and the lambda detection range was set between 415 and 691nm, and intervals set at 8.9nm for the individual detection channels. This permitted the coverage of the whole spectrum with the 32 detection channels. The images were saved in .lsm file format and analyzed with a custom plug-in compatible with Fiji/ImageJ, as described later. Selection of regions of interest was done using ImageJ and the quantification of the GP index from these regions was done as it is explained later.

### D. Quantification of plasma membrane order in plate reader

Measurements of the plasma membrane stiffness were carried out on live NHEKs monolayers, employing either a fluorometer (CLARIOstar, BMG LABTECH, Ortenberg, Germany) or optical microscopy using spectral imaging on a Zeiss 780 inverted microscope. We used an environmental polarity sensitive probe, 6-dodecanoyl-2-dimethylamino naphthalene (Laurdan; Sigma Aldrich). Briefly, for the fluorimetric assessment by the plate reader, cells were seeded out on a 96-well glass-bottom plate (Greiner, Stonehouse, UK). Cell membrane labelling with Laurdan was obtained by 5 to 10min incubation with 50µL of a solution in dimethyl sulfoxide (DMSO, Sigma Aldrich) and Phosphate Buffer Saline pH 7.4 (PBS, Sigma Aldrich), DMSO/PBS 1:3 v/v, at a final Laurdan concentration of 0.5µM at room temperature; afterwards, cells were washed with PBS. Fluorescence emission of Laurdan was exciting at 374nm and recorded over its whole spectrum from 405 to 600nm. The intensity of emission wavelengths at 440±10nm and 490±10nm was used to obtain the GP values. The experiments were done in triplicates employing 3 different cell batches; the fluorescence values observed were an average out of minimum 25 flashes. Additionally, for imaging purposes we added phenol-free L-15 cell media (Leibovitz, Lonza). Samples were prepared and imaged on an 8-well glass-bottom chamber 1.5 (Ibidi, Planegg / Martinsried, Germany).

### E. Generalized polarization index (GP)

Calculation of GP value was carried out as:

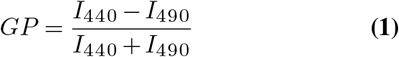

where I_440_ and I_490_ correspond to the emission intensities of Laurdan at 440 and 490nm respectively using 380nm excitation wavelength. Values of GP vary from 1 to −1, where higher numbers reflect lower fluidity or higher lateral lipid order, whereas lower numbers indicate increasing fluidity. The GP measurements on live keratinocyte monolayers at different calcium concentrations (from [Ca^2+^] = 0.06 to 5mM) were on one hand performed on a micro plate fluorescence reader (CLARIOstar, BMG LABTECH), with Laurdan fluorescence emission excited at 374nm and recorded from 405 to 600nm. The emission intensities at 440 and 490 ± 10nm were used to obtain the GP values according to the above equation2. On the other hand, confocal spectral imaging on live NHEKs at calcium concentrations [Ca^2+^] = 0.06 and 1.5mM was performed on a Zeiss LSM 780 confocal microscope equipped with a 32-channel GaAsP detector array. Fluorescence of Laurdan was excited at 405nm and detected between 415 and 691nm. The images were then analysed using a custom plug-in compatible with Fiji/ImageJ, as previously described (35) using a gamma variate fit of the spectra. A frequency histogram of the GP values was generated in Origin Pro (Oregon, USA), which disclosed two populations, one with high GP values representing the plasma membrane fraction and one with low GP values revealing the cytosolic fraction (e.g. from organelles). A fit of a double Gaussian distribution to the distribution allowed determining average GP values as well as standard deviations of both populations. In contrast, the plate reader measurements gave an average value over both populations. For both plate reader and spectral imaging n=3 biological replicate were performed, while from each keratinocyte monolayer a minimum of 25 plate reader recordings or 15 confocal images (spectral images) were acquired and analysed.

For the calculation of the GP index employing spectral imaging and rendering a GP map, we used a purposed-coded plug-in published elsewhere (35) and available at https://github.com/dwaithe/GP-plugin). Briefly, the plugin has in-built the GP general polarization formula and the discrete intensity values around the peak maxima (i.e., 440, and 490nm) are obtained from a fitting to the spectrum obtained per pixel and extracting the values around these peak maxima from the nearest wavelength intervals. This methodology produces a spatial GP map representing the GP value for each pixel of the image. The outputs from these pixels are then exported into OriginPro (OriginLab, Oregon, USA) to produce the normalized frequency counts histograms per GP value. Every image pseudo-color representation has a corresponding custom look-up-table (LUT) matching the highest and lowest GP value obtained from the data.

## Results

### F. Membrane order assessment in live human epidermal keratinocytes at single-cell level

Previously, we extensively characterised the coordinated role of actin scaffolds and filaggrin granule formation during differentiation of normal human epidermal keratinocytes (NHEKs) (9). Specifically, we monitored filaggrin and actin at different calcium concentrations and described and quantified the distribution of these supramolecular assemblies in cornification. We noticed that membrane molecular packing was following an interesting condensation pattern. Therefore, we decided to further investigate this and develop a fast, robust and easy method to assess cornification development by quantifying lipid order in NHEKs. As a control, we first characterised the internal molecular process by observing the filaggrin localisation during calcium switch. Figure 2A shows a dramatic increase of filaggrin-positive granule accumulation in the cytosol and around the nucleus with higher calcium concentration. In order to characterise the lipid lateral packing properties in NHEK membranes, we directly labelled undifferentiated cells grown at low calcium level ([Ca^2+^]=0.06mM) with Laurdan. Next, we carried out spectral imaging by confocal microscopy to address changes in membrane stiffness reported by the quantification of the laurdan general polarization (GP) index. This solvatochromic probe reports on the extent of water relaxation processes in its direct proximity in local molecular environment (Figure 2B-D). An additional advantage of Laurdan is that this dye only fluoresces when incorporated into a membrane environment, hence the GP index is unaffected by the minimal amount of Laurdan which is not associated to membranes. In cells, it is known that the overall internal organelle membrane composition differs from the plasma membrane (22). Plasma membrane lipids have been reported to be more tightly packed and this is clearly demonstrated in Figure 2E GP pseudo-color map, where the plasma membrane display a redder color, which is indicative of higher degree of condensation or more densely packed membranes in comparison to intracellular membranes. Since our aim for this study was to develop a high throughput method to report on keratinocyte differentiation, we wanted to ensure that the overall quantification of the GP index would unambiguously report on the degree of cellular lipid lateral condensation, in a similar fashion as the quantification performed at the plasma membrane only. For this purpose, we compared the GP index population distribution and gaussian fits of a whole cell (Figure 2E-1) with the GP of the internal membranes (Figure 2E-2) and with the GP of the plasma membrane (Figure 2E-3). The distribution of GP values of the whole cell yielded two distinct gaussians: one with lower GP values, mostly distributed in the internal organelles and reflecting more fluid membranes (see Figure 2E in blue-green color; GP=-0.09); and the other with higher values, mainly localised at the plasma membrane, indicating higher degree of lateral packing (Figure 2E in yellow-orange color; GP=0.26). The values for the lower and higher GPs obtained from the whole cell, did not substantially differ from the lower GP values of the cytosolic membrane component and the higher of the plasma membrane component. This indicates that resolving the GP index distribution in cell membranes is sufficient to successfully characterize the lipid order heterogeneity.

### G. Laurdan general polarization index increases during in vitro keratinocyte differentiation

Next, we tested whether confocal spectral imaging would show membrane order changes in live NHEKs during keratinocyte differentiation at a single cell level, in our in vitro model. For this purpose, we employed a well-known and frequently used “calcium switch” assay, in which keratinocytes withdraw from the cell cycle and enter terminal differentiation when exposed to a rapid increase of calcium level in the media. Since a very steep calcium gradient in the epidermis is thought to be the main factor promoting differentiation and stratification in vivo (42–44), this model directly relates to skin physiology. Initially, we validated our assay by simultaneous determination of the differentiation progression in the cells using proteomics (end point assessment). Here, we decided to use the ratio between the expression of a late differentiation marker (filaggrin) and keratin 14, which is predominantly expressed by undifferentiated keratinocytes of lower epidermal layers, including in the basal layer characterized by high proliferation rates (Figure 3A). The increase of the filaggrin/keratin 14 ratio indicates keratinocytes advancing in their differentiation process. Next, using spectral imaging we investigated whether quantifying the GP index in live NHEKs in our validated differentiation model would yield differences that could be confidently used to asses cornification by means of membrane order quantification. Specifically, we exposed NHEKs to increasing calcium levels, labelled them 24h later with Laurdan and determined GP imaging maps and GP index population distributions. The GP maps in Figure 3B confirmed the overall cellular membranes state, showing a decrease in fluid membranes overall and an increase in more ordered membranes. Moreover, the histograms showed a clear and significant response of the membrane packing readout in both the low and high GP value with increased calcium concentration. For instance, from 0.06mM to 1.5mM Ca^2+^ the whole population distribution is shifted towards higher GP values: from −0.06 to 0.05 and from 0.26 to 0.45 for the low and high GP, respectively. Interestingly, the highest calcium concentration (5mM) shows complete disruption of the cells and their membranes. Noteworthy, this lipid lateral packing order changes correlate with what literature has described as an increase in cellular stiffness upon keratinocyte differentiation. Plasma membrane stiffness and thus GP values increased with calcium concentration throughout the tested range (0.06-5mM). Consequently, GP values hold as a robust readout for determination of differentiation stages of keratinocytes at the single live-cell level.

**Fig. 3.**
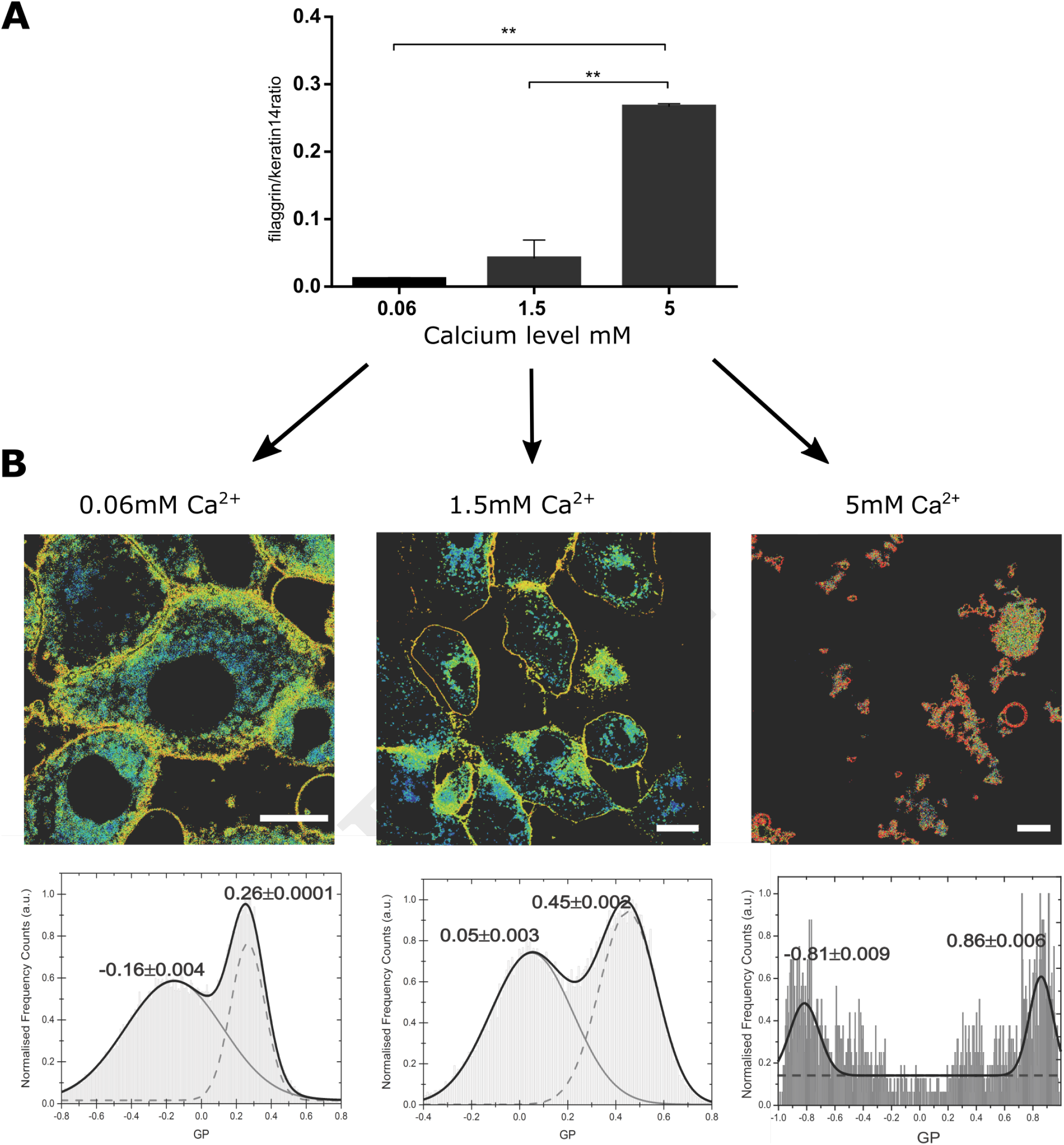
Changes in membrane stiffness during in vitro keratinocyte differentiation by LAURDAN live imaging at the level of single cell. A) Representation of the keratinocyte differentiation advancement by assessment of filaggrin/keratin 14 ratio in intracellular western blot assay in the cells used for LAURDAN assay; representative of 2 separate NHEK batches, assay carried out in duplicates; B) GP images (upper panels) and histograms of GP pixel values (lower panels, from whole image) for increasing calcium levels ([Ca^2 +^]=0.06mM (left), 1.5mM (middle) and 5mM (right)). The histograms were fit by a double-Gaussian (black line, with peak values and their accuracy from the fit stated) to outline the increase in GP and thus stiffness for the plasma membrane fraction (right peaks) and an indifferent change for the cytosolic fraction (left peaks). Representative example from 3 different cell batches, and a minimum of 5 cells per batch; scale bar 5µm.

### H. High throughput GP index quantification assay to determine progression of differentiation in live keratinocytes

Once we demonstrated that the GP index is a suitable reporter on cell membrane order without the need to obtain values exclusively from the plasma membrane, we decided to move further in our aim of developing a cheap and robust methodology to asses keratinocyte differentiation with a high throughput readout and analysis. For this, we tested whether calculating the GP index at the cell-ensemble level, rather than at the single-cell, would yield corresponding results. Hence, we reproduced the previous experimental set up, but employing a plate reader instead of a confocal microscope. This assay provided average GP values from whole cell populations within sub-second time frames. In these experiments we determined GP values representative of entire cell, i.e. calculating an average over both the cytoplasmic and cytosolic membrane fractions. This new approach allowed us to test multiple calcium concentrations faster and in a larger field of view; ultimately, allowing access to a more robust statistical analysis. A caveat to this approach is the fact that the readout is an average of the fluorescence spectra detected from an ensemble of keratinocytes within a cell monolayer. To further test whether this could be a set back to our method, we parallelly ran the samples in a plate under a confocal microscope and in a plate reader (Figure 4A). Notably, using the same cell populations as imaged before on the confocal microscope (the plate format fitted both the microscopy setup and the fluorescent reader) we could still observe a gradual increase of the Laurdan GP values along with the progression of keratinocyte differentiation in our calcium switch model (Figure 4B). The low error bars along with the short measurement times highlight the plate reader assay as a robust high-throughput readout to assess keratinocyte differentiation.

**Fig. 4.**
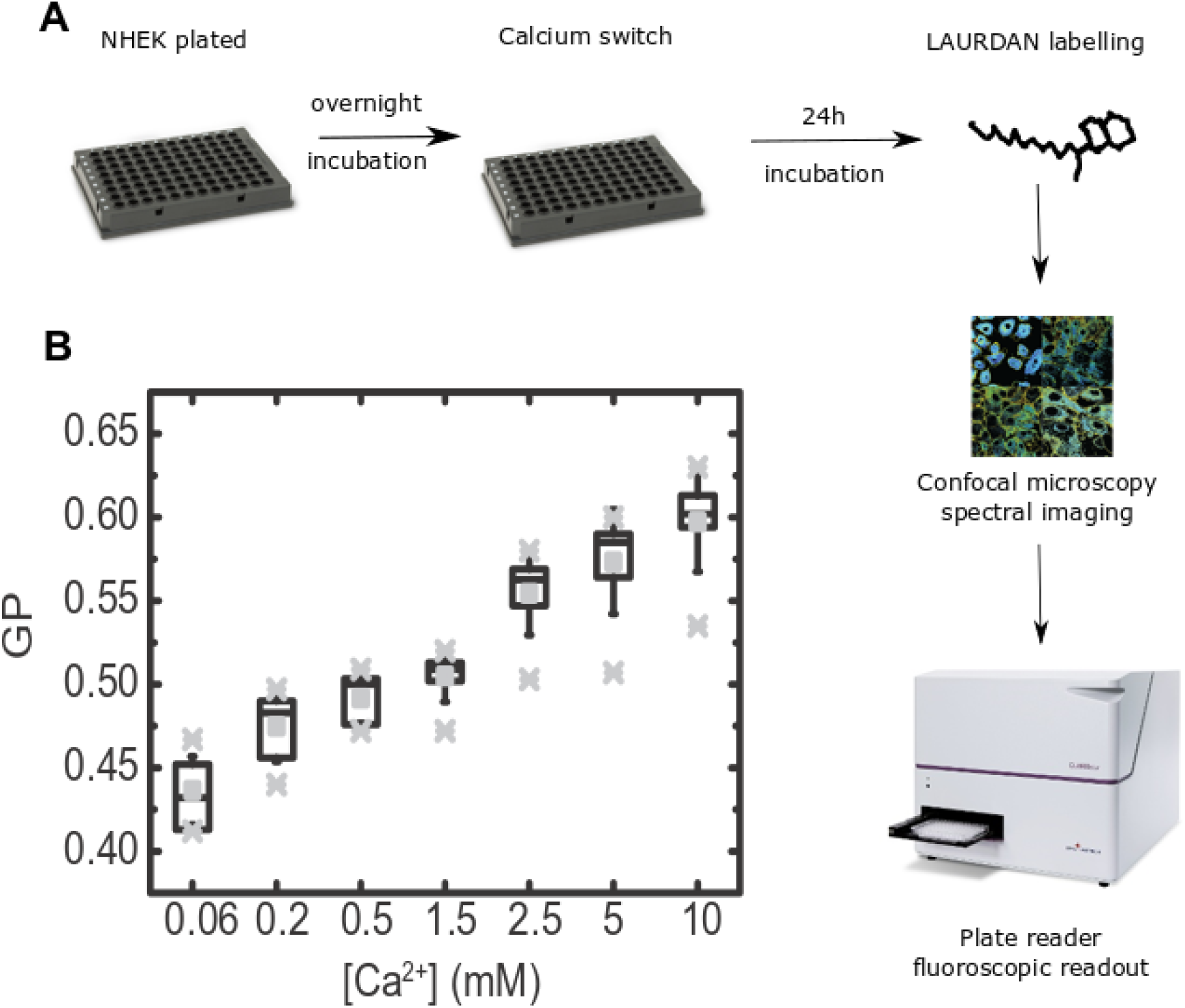
Laurdan General Polarisation (GP) assessment in living keratinocytes at the population level by fluorescent plate reader. A) Workflow diagram: primary keratinocytes were seeded and calcium switch in a range of calcium concentrations ([Ca^2 +^]=0.06mM to [Ca^2 +^]=10mM) at confluence. 24h after cells were LAURDAN-stained live and GP values were obtained by confocal immunofluorescence readout () and fluorescence spectroscopy on a plate reader. B) Whiskers-box plot of the distribution of GP values, with average (middle vertical black lines), the standard deviations (error bars), and the outlier ranges (grey crosses).

## Discussion

Epidermis consists of multiple layers of keratinocytes which represent different stages of progression of terminal differentiation. Successful and complete differentiation is critical for the functional characteristics of the skin, and affects their permeability, antimicrobial and mechanical properties. While the cells gradually progress, their characteristics change dramatically; this results in differential cell reactivity and outcomes in functionally distinct keratinocyte populations. To date there are no specific tools available to trace the progression of keratinocyte differentiation in vitro or to precisely determine differentiation advancement stage of these cells while preserving cell viability; at neither single-cell nor population level. Such an assessment would be beneficial, however, as it would enable the determination of the dependence between the cell reactivity and functional outcomes of their differentiation program advancement. Furthermore, a plate reader format could provide an efficient tool for scaling into a high-throughput assay. Similarly, given that several skin diseases have been found to show a degree of delayed or abnormal epidermal differentiation and resulting insufficiency in the skin barrier function (1, 45), there is an unmet need for robust and validated techniques which could be used to propose new diagnostic and therapeutic options Our study provides cheap and effective methodology to assess keratinocyte differentiation and has implications to both basic and applied research, e.g. in drug testing.

Lipid membrane remodelling and resulting changes in polarity can be employed to measure the degree of condensation of membranes by utilizing fluorescent lipophilic probes such as Laurdan, whose emission spectra are solvent-polarity dependence report on lipid lateral packing. The readout of the Laurdan fluorescence and GP index can be used to provide information regarding membrane heterogeneity, which allows for the estimation of membrane stiffness and fluidity. Furthermore, lack of apparent Laurdan toxicity and the subsecond readout time of the plate reader allowed us to study changes in keratinocyte stiffness and progression of differentiation over time.

In this study we demonstrated that Laurdan general polarization (GP) index can be used as an effective tool to determine keratinocyte differentiation advancement and progression in live keratinocytes at both single-cell and population level, within a cell monolayer and with an option for high-throughput assessment. Laurdan has been previously used in experiments with porcine skin to determine penetration of liposomes (46). Assessment of Laurdan emission in these studies showed a decrease of the GP value with the increased skin depth, from stratum corneum through to the dermis, including a progressive decrease at the level of the epidermis; this is in line with data presented here.

In summary, we designed and validated a Laurdan-based assay which enables for a precise determination of differentiation advancement stage of living keratinocytes, by assessment of changes in membrane stiffness. This in vitro assay, which, due to the lack of apparent cytotoxicity, may also be used to trace differentiation over time, either at the level of single cell or scaled for high throughput measurements.

## Author Contribution

D.G-O and J.B.d.l.S. designed and performed the experiments, carried out the data processing and analysis, discussed, prepared and wrote the manuscript.

C.E. and G.O. supervised the work, helped with analysis, discussed, prepared and wrote the manuscript.

## Acknowledgements

D.G-O. and G.S.O. are grateful for support from the Medical Research Council (UK) and the NIHR Oxford Biomedical Research Centre. D.G.-O. would like to acknowledge the support of British Skin Foundation, European Union’s Horizon 2020 research and innovation programme under the Marie Skłodowska-Curie grant agreement No. 665778, as a part of National Science Centre POLONEZ Fellowship (UMO-2016/23/P/NZ6/04056) and First TEAM programme of the Foundation for Polish Science co-financed by the European Union under the European Regional Development Fund (POIR.04.04.00-00-21FA/16-00). C.E. acknowledges support from the Wolfson Imaging Centre Oxford (Christoffer Lagerholm), the Wolfson Foundation (18272), the Medical Research Council (MCUU12010/unit programmes G0902418 and MCUU12025), MRC/BBSRC/ESPRC (grant number MR/K01577X/1), the Wellcome Trust (104924/14/Z/14, 100262Z/12/Z, 098274/Z/12/Z and Strategic Award 091911 (107457/Z/15/Z, Advanced Micron Bioimaging Unit)), the Deutsche Forschungsgemeinschaft (Collaborative Research Center 1278, Research unit 1905, Jena Excellence Cluster “Balance of the Microverse”), and internal University of Oxford funding (EPA Cephalosporin Fund and John Fell Fund). J.B.d.l.S. acknowledges support from a Marie Curie Career Integration Grant (PCIG13-GA-2013-618914).

